# Optimizing CRISPR/Cas9 System to Precisely Model Plasminogen Activator Inhibitor-1 Point Mutations in Mice

**DOI:** 10.1101/254805

**Authors:** Yang Liu, Thomas L. Saunders, Thomas Sisson, Robert Blackburn, David S. Ginsberg, Duane Day

## Abstract

CRISPR/Cas9 has become a powerful genome editing tool in recent years. CRISPR/Cas9 can be utilized to not only efficiently generate knock out models in various organisms, but also to precisely model human disease or variants to study gene function and develop therapies. However, the latter remains challenging because of low knock-in (KI) efficiency. In this study, precise gene editing modeling plasminogen activator inhibitor-1 (PAI-1) ‐tissue plasminogen activator (tPA) binding deficiency and PAI-1-vitronectin binding deficiency were generated respectively in mice. Optimization of single guide RNAs (sgRNA) and repair templates, and utilization of restriction fragment length polymorphism (RFLP) to detect KI events are described. Injection of sgRNA/Cas9/single-stranded oligodeoxynucleotide (ssODN) into mouse zygotes resulted in homozygous changes of two silent mutations and changed Arg369>Ala, which abolishes PAI-1 inhibitory activity against tPA. Targeting Arg124 and Gln146 simultaneously involved in vitronectin binding proved to be challenging. However, we successfully generated these relatively distant mutations (23 amino acids apart) seamlessly. Generation of the Arg124 mutation alone was achieved with over 60% efficiency along with the integration of a restriction site, compared to the relatively low double mutation frequency. In summary, our data indicates that the distance between desired mutations and CRISPR-induced double-stranded break (DSB) site is the most critical factor for achieving high efficiency in precise gene modification.

## Introduction

Clustered regularly interspaced short palindromic repeats (CRISPR) and CRISPR associated protein 9 (Cas9) have been widely used in gene editing in the last five years. Cas9 is the most used enzyme among all identified Cas proteins, serving as molecular scissors to make double-stranded breaks (DSB) in chromosomes specifically under the direction of CRISPR RNA complimentary to target DNA. In contrast to the tedious construction of zinc finger nucleases (ZFN) and Transcription Activator Like Effector Nuclease (TALEN), CRISPR/Cas9 is relatively simple to design and construct, therefore has become the most popular genome editing tool in basic research and gene therapy across biology, agriculture and medicine.

CRISPR technology has the potential to correct genetic disorders in patient cells or human embryos and provide new therapies and cures for devastating diseases [1]. However, challenges involving specificity and efficiency of gene replacement still exist. Optimization of these factors will enable translation of genome editing technology from laboratory to clinical applications.

Animal models generated by CRISPR/Cas9 system facilitate loss of function and gain of function of preclinical genetic studies. Highly efficient correction depends on precise editing following DSB created by CRISPR/Cas9 in specific genomic loci. To introduce point mutations, small tags or precise deletions in the genome, a repair template or donor is needed. ssODN has been confirmed to be more efficient than other kinds of template such as vector or linearized dsDNA [2, 3]. Different from homology directed repair mediated by double stranded DNA, single stranded template repair mechanism is still not fully understood. Liang et al. [4] proposed a model involving cellular repair upon DSB formation. In the presence of DSB, cellular nuclease generates 3’ overhang to which complimentary ssODN can anneal. Cellular polymerase can then extend 3’ to form a complete strand containing desired mutation. Junctions are repaired by endonuclease and ligase afterwards.

In this study, multiple PAI-1 point mutations were generated with CRISPR/Cas9 reagents together with repair templates in mice using optimized techniques. Strategies for guide selection, optimization and donor design are described. Homozygous changes were observed in all KI mice when the distance between the DSB and the KI is less than 2 bp, while the efficiency of KI dropped greatly when the distance increases to 69 bp but is still feasible. Our data suggested that the distance of mutation to target DSB site is the most critical factor when making small precise mutations in mouse genome.

## Materials and Methods

### Ethics Statement

All microinjection procedures experiments using mice were approved by the Institutional Committee on the Use and Care of Animals for the University of Michigan (protocol number PRO00005913). Mice were housed in specific pathogen-free conditions with ad libitum access to food and water.

### Materials

Cas9 mRNA was purchased from PNABio, Ens-Cas9 Nuclease, T7 High Scribe, NcoI, XbaI, Phusion polymerase, PCR cloning kit were purchased from New England Biolabs. KOD hot start polymerase was purchased from Novagen. RNA clean up kit was purchased from Zymo Research. DNA oligonucleotides used for gRNA synthesis, PCR primers, Invitrogen Maxiscript^TM^ T7 kit were purchased from Thermo Fisher Scientific. ssODNs were purchased from Integrated DNA Technologies (IDT). HB101 Competent cells were from Molecular Innovations.

### sgRNA design and synthesis

Serpine1 genomic sequence was obtained from Ensembl.org. Several online sgRNA design tools were used to screen guides including Broad Institute, www.deskgen.com and http://crispor.tefor.net. On target and off target scores were calculated using published methods [5,6]. The principle is to have maximum on target score and minimum off target sites, close to targeted nucleotides, ideally within 10 bp. Other considerations for choosing guides included targeted nucleotides serving as PAM or PAM proximal, which would prevent Cas9 cleavage after the desired point mutations were introduced. Target specific oligos and universal oligos (sgRNA scaffold) were ordered at IDT (S1 Table) and assembled in a PCR reaction to generate the DNA templates for in vitro transcription of the sgRNA. In brief, the PCR reaction included 100 nM of universal oligo 1, 1 μM of universal oligos 2 and 3, 12 nM of target specific oligos, Phusion polymerase, dNTP and HF buffer supplied in the Phusion kit. PCR was performed using the following cycling conditions: 98°C for 10 sec, 35 cycles of 98°C for 10 sec, 55°C for 15 sec, 72°C for 15 sec, and 72°C for 1 min. PCR products were directly *in vitro* transcribed without further purification using Invitrogen Maxiscript ^TM^ T7 kit or T7 Highscribe (NEB), and resulting RNA was purified using Zymo RNA Clean & Concentrator ^TM^ -25 clean kit, and RNA concentration was determined by BioSpec-nano (Shimadzu Biotech).

### Donor design

Repair template was designed to introduce desired point mutations in specific genomic locus. ssODN serving as repair template was chemically synthesized. ssODN was complementary to the “no PAM” strand. ssODN contained asymmetric homology arms. The left homology arm was ∼36-nucleotide proximal to PAM and right homology arm was ∼ 90-nucleotide distal to PAM [7]. Reference genome sequence and known variants obtained from Ensembl were used as a guideline to exclude single-nucleotide polymorphism (SNP)s in repair template. Target region was sequenced to identify unknown SNPs that would reduce the efficiency of repair. A silent mutation was introduced to generate restriction enzyme cleavage site to ease genotyping of modified animals.

### *In vitro* cleavage assay

Cleavage efficiency was measured *in vitro* using Cas9 protein according to the manufacture’s instruction. Briefly, 100 ng *in vitro* transcribed gRNA and 1 μM Cas9 nuclease were incubated at room temperature for 10 min. 60-100 ng substrate that was PCR amplified of target region and purified was added to Cas9/gRNA complex, and incubated at 37°C for 1 hour. Reaction products were analyzed on 2% agarose gel and gel bands intensity was quantified using Image J [8]. Percentage cleavage was calculated by dividing the intensity of sample band by control band (substrate only).

### Microinjection

Zygote preparation and manipulation, microinjection and embryo transfer to pseudopregnant females followed standard methods [9]. To prepare microinjection solution, 100-200 ng/μL Cas9 nuclease, 50-100 ng/μL gRNA, and 100 ng/μL ssODN were mixed in TE buffer (pH7.4), incubated at room temperature for 10 min, and centrifuged at 13000 rpm for 5 min [10]. Supernatant was stored at - 80°C or on ice for microinjection. Pronuclear alone or both pronuclear and cytoplasmic injected zygotes were transferred to pseudopregnant females at 2-cell stage. Microinjection was performed at the Transgenic Core of University of Michigan. Mice were housed in ventilated racks with automatic watering and ventilated cages. Zygotes for microinjection were obtained by mating C57BL/6J (Jackson Stock Number 000664) or B6SJLF1 (Jackson Stock Number 100012) female mice with B6SJLF1 males. G0 founders were backcrossed with C57BL/6J mice to establish lines.

### Genotyping by PCR-RFLP

Moue tail tip biopsies were collected at 2-week of age, incubated in buffer (100 mM Tris, 5 mM EDTA, 200 mM NaCl, 0.2% SDS, pH8.0) with Proteinase K (0.4 mg/mL) at 55°C overnight. Genomic DNA was isolated by isopropanol precipitation followed by washing with 70% ethanol. PCR primers were designed using Primer 3 [11] and are listed in S1 Table. PCR followed standard conditions, and products were digested using restriction enzyme at 37°C for 30 min and examined on 2% agarose gel. Those showing cleaved bands on the gel were sent for Sanger sequencing directly, or cloned into a vector using PCR cloning kit (NEB). 8-24 clones were screened and RFLP positive clones were sent for Sanger sequencing.

## Results and discussion

### Homozygous change close to DSB were generated in all KI founder mice

PAI-1 is a serine protease inhibitor that plays a key role in the regulation of fibrinolysis. PAI-1 reactive bond residues Arg369 and Met370 are critical to inhibitory activity against tPA [12]. To disrupt the reactive bond, we proposed to mutate Arg369 (CGC) to Ala (GCC) using CRISPR/Cas9 mediated mutagenesis. Genomic sequence of Serpine1 encoding PAI-1 was uploaded to several online tools (see Methods). Based on on-target score, off-target score and location, 4 guides targeting Arg369 were synthesized by *in vitro* transcription, and their on-target activities were tested *in vitro*. The guide demonstrating the highest cleavage efficiency was chosen as the guide to design repair template (Fig 1A). The length of ssODN, sense or antisense, position and structure have been intensively investigated recently [4]. It has been reported that asymmetric ssODN improved targeted integration efficiency [7]. In this study, all the repair templates were asymmetric. The repair template for Arg (R) > Ala (A) comprised 127 nucleotides, with 36 nucleotides on the PAM-distal side of DSB and 96 nucleotides on the proximal side. NcoI site was generated simultaneously when CGC was changed to GCC with no need to make additional changes to create restriction site. To prevent unwanted cleavage after desired editing was completed, two silent mutations were made proximal to PAM: Thr373 ACG>ACT, Glu374 GAG>GAA. Phosphorothioate modified ssODN has been shown enhanced efficiency and greater flexibility in cultured cells and rodents [12]. ssODN was ordered at IDT with phosphorothioate modifications at 3’ and 5’ ends (s3 Table) with no further PAGE purification.

**Fig 1.**
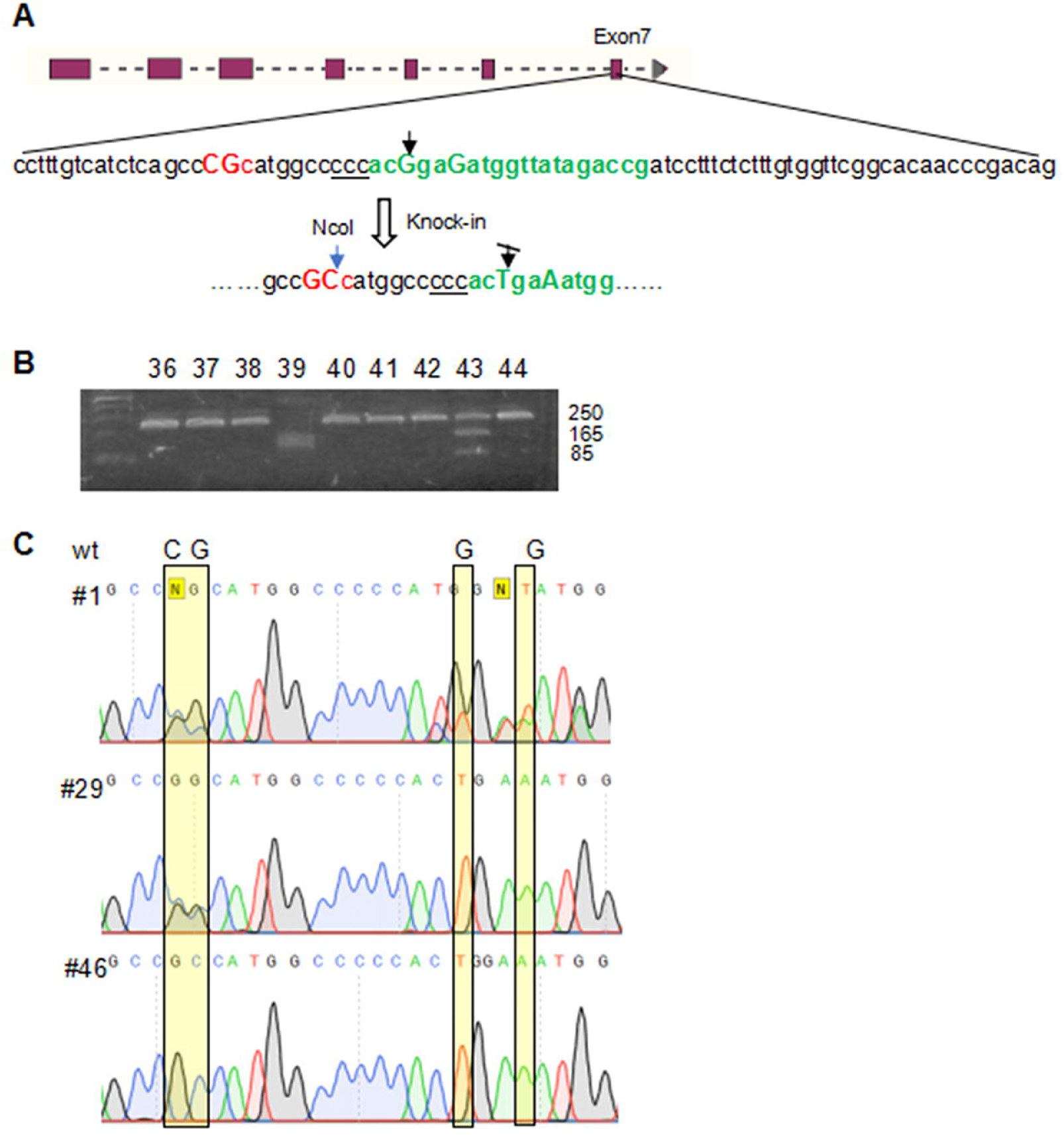
Homozygous changes close to DSB were generated in all KI founder mice. **A.** Partial Exon 7 sequence shows the nucleotides to be changed CG>GC (in red), PAM site (underlined), guide sequence (in green) and two silent mutations (in green, bold). Black arrow indicates Cas9 cleavage site. Blue arrow indicates the NcoI cleavage site after the integration of repair template. **B**. The PAI-1 Arg369 genomic locus was PCR-amplified, followed by NcoI digestion. Representative gel image shows one positive KI (#43) and one large deletion (#39) identified by PCR-RFLP. **C**. Examples of edited animals. Yellow bar indicates four targeted nucleotides. Pup #1 was a mosaic founder containing both KI and KO alleles. Cloning and Sanger sequencing of colony PCR products confirmed the homozygous change of two silent mutations. #29 had the heterozygous change of R>A (CGC>GCC), homozygous changes of two silent mutations. #46 had the homozygous changes of all four targeted nucleotides.

100 ng/μL Cas9 mRNA, 100 ng/μL ssODN and 100 ng/μL gRNA were injected into mouse zygote pronuclei. Total 177 zygotes were injected, and 108 2-cell zygotes were transferred to pseudopregnant females. 46 pups were born and tailed at 2-week old. Tail DNA was used as a template to amplify target region followed by NcoI digest (Fig 1B).

RFLP assay has been widely used for evaluating editing events [13], and 6 samples (6/46, 13%) were identified as NcoI cleavage positive and PCR products were sent for Sanger sequencing directly, or cloned into a vector and colony PCR products were sequenced. Sequencing results confirmed that all six pups were PAI-1 R>A founders, and 3 being heterozygous, 2 mosaic and 1 homozygous (Fig 1C). Interestingly, these six founders all had homozygous change of two silent point mutations. Given that the Arg369 is 13-14 bp away from DSB, while two silent point mutations introduced for blocking Cas9 are 0 and 2 bp away from DSB respectively and Cas9 cleavage site is 3bp adjacent to 5’-PAM, the frequency of R>A mutation (17%) relation to that of the silent mutation (100%) appears to be affected by proximity to the DSB.

### Optimize guide RNA to achieve high on target activity *in vitro*

PAI-1 Arg124 and Gln146 are two key amino acids involved in vitronectin binding. Previous PAI-1 R>A editing data has shown that editing efficiency greatly decreased as the distance to cleavage site increased. Since there are 23 amino acids (69bp) between Arg124 and Gln146, we considered to use two sgRNA targeting Arg124 and Gln146 respectively (Fig 2A). Three guides were chosen around each target site based on distance, on-target and off-target scores. The oligos were ordered and assembled, and guide RNAs were *in vitro* transcribed and screened in an *in vitro* cleavage assay.

**Fig 2.**
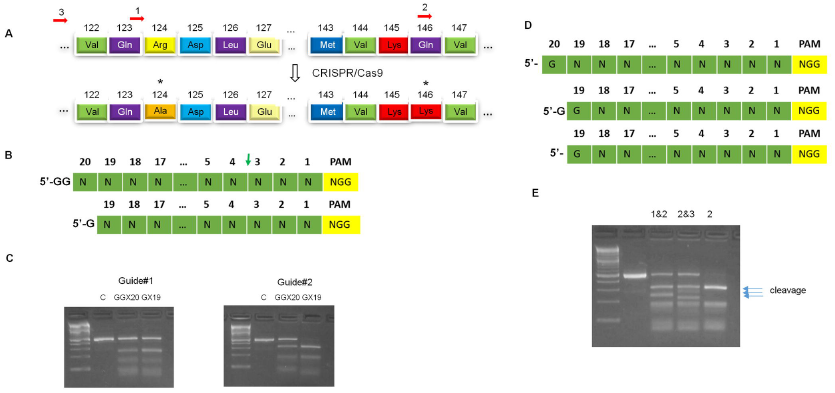
Optimization of guides targeting PAI-1 vitronectin binding reactive residues Arg124 and Gln146. **A.** The two expected mutant amino acids Ala124 and Lys146, which are indicated by asterisk, would abolish vitronectin binding activity. Red arrows indicate three guides targeting Arg124 and Gln146, respectively. **B**. Standard guide sequence GGX20 contains 20 nucleotides adjacent to NGG (PAM), GG is added to 5’ end to initiate transcription *in vitro*. GX19 contains 19 nucleotides plus one G added to 5’ end. Arrow in green indicates Cas9 cut site. **C**. *In vitro* cleavage assay showed that guide 1 in the format of GGX20 and GX19 had no difference in cleavage, but guide 2 in GX19 had significant increased cleavage efficiency compared to GGX20. **D**. Guide sequence in different length had variable cleavage performance *in vitro*. **E**. Comparison of multiplexed guides 1&2 and guides 2&3 versus guide 2 alone in the *in vitro* cleavage assay. Guide 1 which is closer to target was picked together with guide 2 for the repair template design.

Guide 1 and guide 2 were picked respectively for target sites for showing highest cleavage activity. We have observed *in vitro* activity was not always consistent with activity *in vivo* (unpublished data), however, *in vitro* testing allows a quick screen of multiple guides, and is an indication of the quality of CRISPR reagents prior to proceeding to microinjection.

In general, guide sequence contains 20 nt adjacent to PAM (NGG). We routinely added two G to the guide sequence to initiate T7 *in vitro* transcription (Fig 2B). Guide 1 showed only about 50% cleavage in the *in vitro* assay. To achieve higher cleavage activity, we first tested various *in vitro* transcription kits to generate guide RNAs but did not observe significant differences in terms of cleavage *in vitro*.

It has been reported that the addition of extra G added to 5’ guide sequence for initiating *in vitro* transcription reduced editing efficiency [14,15]. Truncated guide RNA (17-18 nt) has been shown to improve specificity without compromising on target cleavage activity [16]. To test the cleavage efficiency of truncated guide RNA with only one additional G, we made new guide 1 with the sequence 5’-Gggatgccatctttgtccag (GX19) in comparison with the previous 5’-GGcggatgccatctttgtccag (GGX20), and similar change to guide 2: 5’-Gcagactatggtgaaacagg (GX19) versus 5’-GGccagactatggtgaaacagg (GGX20). The yield of transcribed RNA *in vitro* decreased when GG was changed to G but was sufficient for *in vitro* assays and microinjections. GGX20 and GX19 guide RNAs were compared in *in vitro* cleavage assay (Fig 2C). Cleavage of substrate mediated by guide 2 increased from about 50% (GGX20) to 90% (GX19), however, guide 1 (GX19) didn’t improve significantly.

We have tested sgRNA in various formats when targeting genes that encode other coagulation proteins. GX19 sgRNA didn’t always increase cleavage activity dramatically but at least retained similar activities as GGX20. Some algorithms set N20 as G or N19-20 as GG default when searching for guides, which may exclude guides that have high on-target activity but N20 is A, T or C. We observed that when N19 is G, the guide X19 performed better than GX19 *in vitro* (Fig 2D). Overall, GX18-19 guides demonstrated equivalent or greater on target activity as compared to GGX20 guides.

In order to develop a guide RNA targeting Arg124 with high on target activity, we tested chemically synthesized guide RNAs in full length (X20, N20≠G) side by side with their counterpart transcribed GX19 RNAs *in vitro*. Synthetic RNAs have been proven to have better purity and do not require an extra G. Interestingly, full length synthetic RNAs didn’t exhibit greater efficiency than GX19 guides (unpublished data), indicating that G at 5’ end might play a role in RNA stability or Cas9 binding, which lead to higher editing efficiency, though further confirmation is needed.

### Multiplexing guides may affect Cas9 binding capacity

Apparently, distance of mutation to DSB plays a critical role in the integration of the repair template. Still, it is hard to predict the editing efficiency either using a low on-target score guide close to the DSB or a high on-target score guide in which the distance will be compromised. We picked a new guide (#3) targeting Arg124 with 74% cleavage activity *in vitro* but 23 bp away from Arg124, and compared it with guide #1 for the synergic cleavage activity combining with guide #2 *in vitro*.

To test cleavage efficiency of multiplexed guides, guides 1 and 2, and guides 3 and 2 were incubated with Cas9 protein, respectively. However, cleavage assay showed efficient cleavage by guide 2 only, indicating the possibility of competitive binding to Cas9 when multiplexing guides. Guides were then incubated with Cas9 individually to form RNA/Cas9 complex prior to combining for *in vitro* cleavage. Results showed successful cleavage by all guides (Fig 2E). Multiplexed guides 1&2 were confirmed to have comparable cleavage activity as guides 3&2, and were chosen as templates for designing donor ssODNs.

### Unwanted mutation from synthesized ssODNs

The mechanism of ssODN as repair template is still not fully understood. When a DSB is formed upon Cas9 mediated cleavage, cellular nuclease generates 3’ overhangs and DNA polymerase then extends 3’ overhangs using ssODN as template which is complementary to 3’ overhang. In this study two ssODNs were synthesized as repair template for the Arg124 and Gln146 mutation respectively with phosphorothioate modification at 1-2 terminal nucleotides at 5’ end and 3 terminal nucleotides at 3’ end. Both contained modification CGg>GCg and Cag>Aag that converts Arg to Ala and Gln to Lys, and silent mutations to create restriction enzyme cut sites, respectively (Fig 3A). Changes (CGg>GCg, Cag>Aag) would block Cas9 recut after edits were completed. SNPs have been known to reduce editing efficiency, however, we were not able to exclude a known A/G SNP which is only 14 bp away from Gln146.

**Fig 3.**
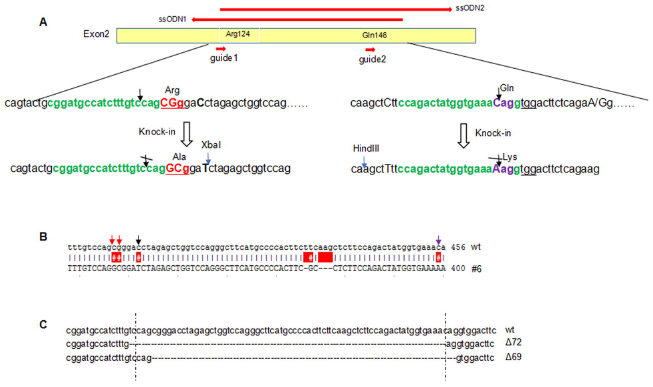
Unwanted mutations were identified when using dual guides/donors. **A.** Schematic showing wild type alleles, targeted alleles, and location of two guides and two ssODNs (red arrow). Donors containing desired mutations were designed and synthesized. Partial exon 2 sequence shows target sequences (in green), PAM (underlined), Arg124 > Ala (in red), Gln146 >Lys (in purple), nucleotides to be mutated (capitalized), Cas9 cut site (black arrow), restriction enzyme digestion site (blue arrow), and a A/G SNP (capitalized). **B.** Cloning and sequencing of PCR products derived from G0 founder pup #6 revealed desired mutations CG>GC (Arg124 > Ala) and C>A (Gln 146 > Lys), and an unwanted 4 bp deletion on exon 2. **C.** Large deletions were identified in G0 pups, demonstrating highly efficient editing mediated by dual guides. Dashed line indicates predicted Cas9 cut site of each target.

Cas9 ribonucleoproteins (RNPs) containing purified Cas9 protein and gRNA have been reported to have advantages over Cas9 mRNA/gRNA for efficient gene editing [17]. Interestingly, there are controversial findings in terms of injecting RNPs into pronuclear or cytoplasm of mouse zygotes to obtain higher editing efficiency [18]. Since mouse zygotes tolerate CRISPR reagents well, we chose to inject into both pronuclear and cytoplasm. Total 100 ng/ μL guide RNAs, 200 ng/ μL Cas9 protein, 100 ng/ μL ssODNs were injected into zygotes. 96 2-cell stage eggs out of 157 injected eggs were implanted, resulting in the birth of 8 pups. PCR-RFLP results showed pup #6 was XbaI cleavage positive and #8 had large deletion. None of these pups was HindIII cleavage positive, indicating ssODN2 was not integrated. Cloning and sequencing of PCR products revealed that #6 had two mutant alleles including a 72 bp deletion, and one allele had the correct changes of CGg>GCg and Cag>Aag but also contained an unwanted 4 bp deletion on Exon2 (Fig 3B). Pup #8 was confirmed to possess a 69 bp deletion by sequencing (Fig 3C). The unwanted 4 bp deletion may have resulted from mutations generated in oligo synthesis. Fig 3C shows the large deletions that were created by two guides, consistent with the observation of gene disruptions using 2-4 guide RNAs targeting key exon for generating knock out animal models [19].

### Double mutations were successfully generated but efficiency varied

Based on previous data, to avoid error-prone nonhomologous end joining (NHEJ) ‐mediated large deletion, we chose to use single gRNA/ ssODN instead of double gRNAs/ ssODNs. Since the guide that targets Arg124 and ssODN1 was confirmed to work *in vivo*, and the known A/G SNP needs to be excluded in repair template, furthermore, some publications have shown single mutation (Arg>Ala) can abolish the binding of PAI-1 to vitronectin [20], guide 1 and ssODN1 were chosen for microinjection. Chemical synthesis and chemical modification of ssODN have been described to be toxic to cultured cells [21]. The ssODN was PAGE purified to increase purity, and minimize toxicity and unwanted mutations.

228 zygotes were injected with 100 ng/ μL guide1 RNA, 100 ng/ μL PAGE purified ssODN1 and 200 ng/ μL Cas9 protein. 178 injected zygotes were transferred to pseudopregnant mice and 61 pups were born (Table 1).

**Table 1.**
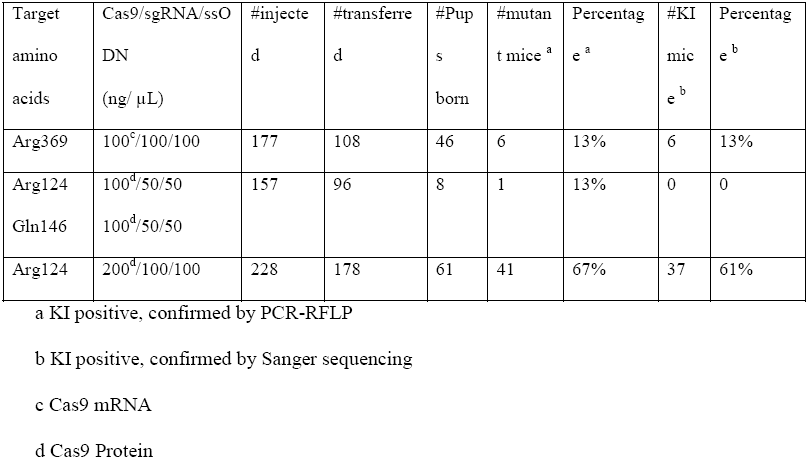
CRISPR/Cas9-mediated Serpine1 targeting in C57BL/6J mice

Tail DNA genotyping revealed that 41 out of 61 pups were KI positive as determined by PCR-RFLP (Fig 4A). PCR products were sent directly for Sanger sequencing except for the 3 samples showing >100 bp deletions in the target region in addition to presence of XbaI cut site, which required further cloning and screening. Of those 20 RFLP negative samples, one (#754) presented a single band on agarose gel with similar size as XbaI cleaved products, and was confirmed to be a biallelic 354 bp deletion by sequencing (Fig 4B), demonstrating that one single guide could generate large indels and should be considered when designing genotyping primers.

**Fig 4.**
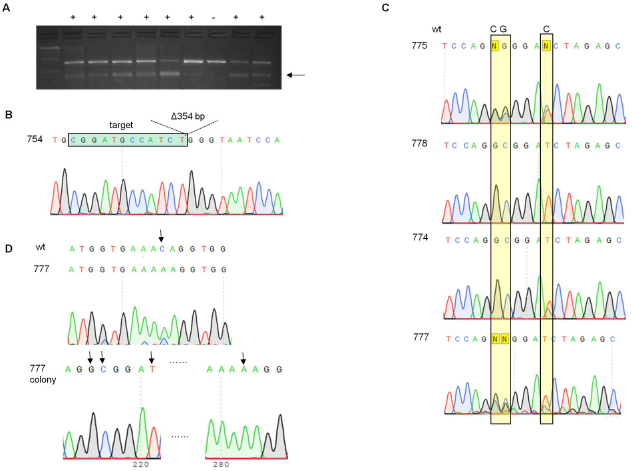
Generation of double-mutation allele using one guide/one ssODN. **A.** Cas9 RNP targeting Arg124 and ssODN were microinjected into mouse zygotes. G0 founder pups derived from the injected zygotes were tailed at 2-week of age and the region flanking Arg124 and Gln146 was PCR amplified, followed by restriction digestion with XbaI. An example of genotyping gel image showed PCR-RFLP results of nine pups on a 2% agarose gel. +, PCRRFLP positive, -, PCR-RFLP negative. The expected size of PCR amplicon is 763 bp, XbaI digested products are 394 bp and 369 bp. **B.** Chromatogram shows a biallelic 354 bp deletion identified in amplified PCR product derived from G0 pup #754. **C.** Representative chromatograms showed various modifications in target region. Yellow bars indicate sites to be mutated containing CG>GC (Arg>Ala) and silent point mutation C>T. G0 pups #775 and 777 had heterozygous changes of all three nucleotides; #778 had homozygous changes of both CG>GC and C>T; #774 had homozygous change of CG>GC but heterozygous change of C>T. **D.** Sequencing chromatogram showed that #777 had heterozygous change of C>A (Gln>Lys). Subsequent cloning, colony screening and Sanger sequencing confirmed that #777 contained double mutations Arg>Ala and Gln>Lys. Arrow in black indicates the nucleotides that have been modified by CRISPR mediated KI.

Sequencing results of 38 RFLP positive samples revealed that one sample that had a 16 bp deletion and an introduced XbaI cut site was a false positive for knock-in. The rest 37 were all confirmed to have the change of Arg>Ala and silent mutation C>T. Arg124 was 3 bp away from the predicted Cas9 cleavage site, resulting in 61% success rate. Of those 37 pups, 11 (30%) had biallelic change of Arg>Ala, and 8 out of those 11 (8/37, 22%) had biallelic change of C>T, which is 8 bp away from cleavage site. Fig 4C illustrates 3 representative gene editing events: #775 and #777 had heterozygous change of Arg>Ala and C>T, #778 had homozygous change of Arg>Ala and C>T, and #774 had homozygous change of Arg>Ala but heterozygous change of C>T. Sequencing results revealed that only #777 had the Gln>Lys mutation as confirmed by cloning, colony PCR and sequencing, as shown in Fig 4D, while the others remained unedited. Efficiency of Arg>Ala could be underestimated by PCR-RFLP, since the XbaI site was 5 bp further from the predicted DSB, therefore XbaI cleavage positive could be a good indication for presence of double mutations (Arg>Ala and Gln>Lys). While Gln146 was 69 bp away from predicted Cas9 cleavage site, unsurprisingly the efficiency dropped down to 2%. Even though the efficiency is low, our data indicated that knock-in events could happen beyond the commonly recommended distance‐‐‐within 10 or 20 bp from the DSB. It would be interesting to test the efficiency of nucleotide changes >30 bp away from each side of DSB, unfortunately a guide targeting in the middle of Arg146 and Gln146 with high activity scores was not available in this case.

Our data suggested that the distance of the change to DSB is the most critical factor for achieving highly efficient precision editing. It is often difficult to select a perfect guide which fulfills all expectations, a compromise must often be made between selecting a guide creating DSB very close to the change but less efficient in cleavage, and one that is further away but with high on target activity. The best scenario will be to choose a guide as close as possible to the desired mutations having reasonable on-target and off-target scores. A recent report has found that single stranded template repair is independent of NHEJ events, and repair relies on Fanconi Anemia pathway but not RAD51-related homology directed repair [22]. The association of ssODN mediated repair and NHEJ event has not been fully characterized. Our data suggests that priority should be given to distance to the change, off-target effects, and on-target score from high to low when choosing guides in silico prior to screening for *in vitro* or *in vivo* cleavage activity.

## Conclusions

Our study has shown PAI-1 single and multiple point mutations were generated in mice by CRIPSR/Cas9 mediated genome editing. We have optimized guide RNA regarding the length and position of G. In our hand, GX19 guide RNA showed the highest cleavage activity *in vitro*. For the design of repair templates, we have considered the length of donor, single vs. dual donors, position of restriction sites, and the existence of known SNPs. We recommend phosphorothioate modification and PAGE purification when using ssODN as a repair template. We didn’t compare symmetric and asymmetric donors but asymmetric donor proved to work well *in vivo* in this study, even when the desired mutation was 69 bp away from DSB. Our data showed that the distance of the change to predicted Cas9 cleavage site correlated with precise editing efficiency, indicating distance is the key player in designing precision gene modifications.

## Acknowledgments

We acknowledged Dr. Jordan Shavit, Steven Grzegorski in the Medical School of University of Michigan, staff members in the Transgenic Core of University of Michigan, Keith Child, Brian Maddaford, Tarek Gharib and other staff in Molecular Innovations for providing technical support and inputs on the experimental design and manuscript.

## References

1. Ma H, Marti-Gutierrez N, Park S-W, Wu J, Lee Y, Suzuki K, et al. Correction of a pathogenic gene mutation in human embryos. Nature. 2017. 548(7668):413–9.

2. Yoshimi K, Kunihiro Y, Kaneko T, Nagahora H, Voigt B, Mashimo T. ssODN-mediated knock-in with CRISPR-Cas for large genomic regions in zygotes. Nat Commun. 2016. 7:10431.

3. Ma X, Chen C, Veevers J, Zhou X, Ross RS, Feng W, et al. CRISPR/Cas9-mediated gene manipulation to create single-amino-acid-substituted and floxed mice with a cloning-free method. Sci Rep. 2017. 7:42244.

4. Liang X, Potter J, Kumar S, Ravinder N, Chestnut J. Enhanced CRISPR/Cas9-mediated precise genome editing by improved design and delivery of gRNA, Cas9 nuclease, and donor DNA. J Biotechnol. 2017. 241:136–46.

5. Doench JG, Fusi N, Sullender M, Hegde M, Vaimberg EW, Donovan KF, et al. Optimized sgRNA design to maximize activity and minimize off-target effects of CRISPR-Cas9. Nat Biotechnol. 2016. 34(2):184–91.

6. Hsu PD, Scott DA, Weinstein JA, Ran FA, Konermann S, Agarwala V, et al. DNA targeting specificity of RNA-guided Cas9 nucleases. Nat Biotechnol. 2013. 31(9):827–32.

7. Richardson CD, Ray GJ, DeWitt MA, Curie GL, Corn JE. Enhancing homology-directed genome editing by catalytically active and inactive CRISPR-Cas9 using asymmetric donor DNA. Nat Biotechnol. 2016. 34(3):339–44.

8. Schneider CA, Rasband WS, Eliceiri KW. NIH Image to ImageJ: 25 years of image analysis. Nat Methods. 2012. 9(7):71–5.

9. Ittner LM, Götz J. Pronuclear injection for the production of transgenic mice. Nat Protoc. 2007. 2(5):1206–15.

10. Qin W, Kutny PM, Maser RS, Dion SL, Lamont JD, Zhang Y, et al. Generating Mouse Models Using CRISPR-Cas9-Mediated Genome Editing. In: Current Protocols in Mouse Biology. Hoboken, NJ, USA: John Wiley & Sons, Inc.; 2016. p. 39–66.

11. Untergasser A, Cutcutache I, Koressaar T, Ye J, Faircloth BC, Remm M, et al. Primer3— new capabilities and interfaces. Nucleic Acids Res. 2012. 40(15):e115.

12. Renaud J-B, Boix C, Charpentier M, De Cian A, Cochennec J, Duvernois-Berthet E, et al. Improved Genome Editing Efficiency and Flexibility Using Modified Oligonucleotides with TALEN and CRISPR-Cas9 Nucleases. Cell Rep. 2016. 14(9):2263–72.

13. Jacobi AM, Rettig GR, Turk R, Collingwood MA, Zeiner SA, Quadros RM, et al. Simplified CRISPR tools for efficient genome editing and streamlined protocols for their delivery into mammalian cells and mouse zygotes. Methods. 2017. 121–122:16–28.

14. Tycko J, Myer VE, Hsu PD. Methods for Optimizing CRISPR-Cas9 Genome Editing Specificity. Mol Cell. 2016. 63(3):355–70.

15. Kulcsár PI, Tálas A, Huszár K, Ligeti Z, Tóth E, Weinhardt N, et al. Crossing enhanced and high fidelity SpCas9 nucleases to optimize specificity and cleavage. Genome Biol. 2017. 18(1):190.

16. Fu Y, Sander JD, Reyon D, Cascio VM, Joung JK. Improving CRISPR-Cas nuclease specificity using truncated guide RNAs. Nat Biotechnol. 2014. 32(3):279–84.

17. DeWitt M, Corn JE, Carroll D. Genome editing via delivery of Cas9 ribonucleoprotein. Methods. 2017. 121-122:9-15.

18. Raveux A, Vandormael-Pournin S, Cohen-Tannoudji M. Optimization of the production of knock-in alleles by CRISPR/Cas9 microinjection into the mouse zygote. Sci Rep. 2017. 7:42661.

19. Zuo E, Cai Y-J, Li K, Wei Y, Wang B-A, Sun Y, et al. One-step generation of complete gene knockout mice and monkeys by CRISPR/Cas9-mediated gene editing with multiple sgRNAs. Cell Res. 2017. 27(7):933–945.

20. Jensen JK, Durand MK V, Skeldal S, Dupont DM, Bødker JS, Wind T, et al. Construction of a plasminogen activator inhibitor-1 variant without measurable affinity to vitronectin but otherwise normal. FEBS Lett. 2004. 556(1–3):175–9.

21. Rios X, Briggs AW, Christodoulou D, Gorham JM, Seidman JG, Church GM. Stable gene targeting in human cells using single-strand oligonucleotides with modified bases. Korolev S, editor. PLoS One. 2012. 7(5):e36697.

22. Richardson CD, Kazane KR, Feng SJ, Bray NL, Schaefer AJ, Floor S, et al. CRISPRCas9 Genome Editing In Human Cells Works Via The Fanconi Anemia Pathway. Preprint. Available from: https://www.biorxiv.org/content/early/2017/05/09/136028

